# Summer drought decreases the predictability of local extinctions in a butterfly metapopulation

**DOI:** 10.1101/863795

**Authors:** Erik van Bergen, Tad Dallas, Michelle F. DiLeo, Aapo Kahilainen, Anniina L. K. Mattila, Miska Luoto, Marjo Saastamoinen

## Abstract

The ecological impacts of extreme climatic events on population dynamics and/or community composition are profound and predominantly negative. Here, using extensive data of an ecological model system, we test whether predictions from ecological models remain robust when environmental conditions are outside the bounds of observation. First, we report a 10-fold demographic decline of the Glanville fritillary butterfly metapopulation on the Åland islands (Finland). Next, using climatic and satellite data we show that the summer of 2018 was an anomaly in terms of water balance and vegetation productivity indices across the habitats of the butterfly, and demonstrate that population growth rates are strongly associated with spatio-temporal variation in climatic water balance. Finally, we demonstrate that covariates that have previously been identified to impact the extinction probability of local populations in this system are less informative when populations are exposed to (severe) drought during the summer months. Our results highlight the unpredictable responses of natural populations to extreme climatic events.

## 1. INTRODUCTION

One of the major challenges in conservation biology is to identify the species and populations that are most vulnerable to extinction. Long-term monitoring of ecological model systems, both at local and global scales, have facilitated conservation objectives by identifying the factors affecting population declines and extinctions (Hanski et al. 1995; Pimm et al. 2014; Dornelas et al. 2019). In addition, ecological models aiming to improve our understanding of population dynamics in temporally varying environments have been employed to shed light on which regions should receive priority for conservation and to predict which species and/or populations are most vulnerable (e.g. Franklin et al. 2014; Oliver et al. 2015). In temporally autocorrelated environments, where conditions tend to change gradually, these predictions may be reliable over short timescales, and can therefore be used to make conservation efforts more effective.

When populations are exposed to conditions that are beyond the normal range, such as in the case of extreme climatic events (ECEs), the factors underlying population dynamics may be less relevant and consequently predictions relying on these factors less reliable. ECEs have increased in recent decades, not only in frequency but also in intensity and duration (Jentsch et al. 2007; Coumou & Rahmstorf 2012; Ummenhofer & Meehl 2017). Recent studies have demonstrated that even a single ECE, such as a flood or a drought, can have profound impacts on natural populations (Bailey & van de Pol 2016; Altwegg et al. 2017; Grant et al. 2017), and can even cause the collapse of an entire ecosystem (Anderson et al. 2017; Harris et al. 2018). For example, a pan-tropical episode of coral bleaching, triggered by a marine heatwave in 2016, eradicated over 60% of the coral communities in the Great Barrier Reef (Hughes et al. 2017).

Studies assessing impacts of ECEs are generally conducted by performing perturbation experiments (e.g. Bokhorst et al. 2008; Pansch et al. 2018) or by opportunistically taking advantage of a rare event (e.g. Smith 2011; Grant et al. 2017). The latter often concentrate on a single population and relatively small spatial scales, or lack long-term ecological monitoring of the system prior to the ECE (but see Thibault & Brown 2008; Campbell-Staton et al. 2017). Here, we report a dramatic demographic decline of the Glanville fritillary butterfly (*Melitaea cinxia*) metapopulation on the Åland archipelago in south-west Finland due to a severe drought in the summer of 2018. This iconic butterfly metapopulation has become an important ecological model system for studying the factors affecting local population dynamics and extinction risks in fragmented landscapes (Hanski 1998; Ovaskainen & Saastamoinen 2018). Due to (ongoing) systematic monitoring, the system provides a unique opportunity to improve our understanding of how large spatially-structured natural populations respond to extreme climatic conditions, and to study how ECEs affect the value of predictive models.

Since 1993, the occupancy and abundance of local populations of the butterflies, across a network of about 4400 dry meadows, has been systematically monitored. Estimates of local population sizes are acquired during autumn when all potential habitat patches are surveyed for the presence of the winter nests (for details see; Ojanen et al. 2013). Despite increasing year-to-year fluctuations (Hanski & Meyke 2005; Tack et al. 2015), the overall size of the metapopulation has remained relatively stable, with about 20% of the available habitat patches being occupied each year. Survey data have been used to demonstrate that the long-term persistence and population dynamics (local extinctions and re-colonizations) of the metapopulation largely depends on the number of available habitat patches, their area, and their spatial configuration (i.e. connectivity; Hanski & Ovaskainen 2000; Dallas et al. 2019). Other key factors impacting local extinctions and population dynamics are the habitat quality (e.g. amount of host plants; Harrison et al. 2011; Schulz et al. 2019) and genetic characteristics of the individuals (Saccheri et al. 1998; Niitepõld & Saastamoinen 2017).

We recently demonstrated that variation in population growth rates is strongly associated with variation in temperature and precipitation across the habitat patch network (Kahilainen et al. 2018). Spatial variability in these climatic conditions contributes to the overall stability of the metapopulation by ensuring that local extinctions are compensated by stable or increased population sizes and colonization rates in other areas of the metapopulation. As the climatic conditions have become more homogeneous across Åland in the last decade, the Glanville fritillary metapopulation has now become more spatially synchronous in its demographic dynamics (Kahilainen et al. 2018), potentially making it more vulnerable to large-scale environmental perturbations (Hanski & Woiwod 1993).

Here, we identify the key anomalies that occurred in the summer of 2018 on the Åland islands and demonstrate that exposure to these extremes was sufficient to drive the regional population declines of the butterfly observed across the metapopulation. Secondly, we explore how the ECE impacts the performance of ecological models by testing whether the covariates that have previously been identified to affect local extinctions in this system remain informative under extreme climatic conditions.

## 2. MATERIAL AND METHODS

### 2.1 Long-term data survey

On the Åland islands, the Glanville fritillary inhabits dry meadows and pastures with at least one of the two larval host plant species, the ribwort plantain (*Plantago lanceolata*) or the spiked speedwell (*Veronica spicata*) present. The entire study region (50 × 70 km) has been mapped for potential habitat patches, with a total of 4,415 patches (September 2015). The population occupancy (presence of a larval nest within a habitat patch) and abundance (total number of larval nests within a habitat patch) is assessed annually when the larvae have entered the overwintering stage within their conspicuous larval nests. All the habitat patches on the archipelago are inspected with the help of approximately 50 field assistants. In the field, the locations of the larval nests are marked and data are stored into the EarthCape database (Ojanen et al. 2013). The occupancy and abundance data comprise raw counts, which is a function of both population size but also of detection probability. Estimates from control surveys conducted before 2018 suggest that the presence of the butterfly may be missed in up to 15% of the occupied patches, with non-detection mainly occurring in very small populations (Hanski et al. 2017).

### 2.2 Climatic and vegetation index data

Normalized Difference Vegetation Index (NDVI) and the Enhanced Vegetation Index (EVI) values were derived from Moderate Resolution Imaging Spectrometer (MODIS, resolution 250 m; Guay et al. 2014). Both NDVI and EVI data are available on a 16-day basis for a 19-year period between 2000 and 2018. Data are derived from bands 1 and 2 of the MODIS on board NASA’s Terra satellite. A time-series of NDVI and EVI observations can be used to examine the dynamics of the growing season, ecosystem health or monitor phenomena such as droughts. EVI is an ‘optimized’ vegetation index designed to enhance the vegetation signal with improved sensitivity in high biomass regions and improved vegetation monitoring through a decoupling of the canopy background signal and a reduction in atmosphere influences. NDVI and EVI values were calculated for each of the 4,415 habitat patches from April to August. Monthly temperature and precipitation values and water balance (precipitation - potential evaporation) are derived from Jomala weather station (60.18° N, 19,99° E, 11 m a.s.l.) database. Climatic water balance was quantified by precipitation minus potential evapotranspiration (PET). PET is the amount of evaporation and transpiration that would occur if a sufficient water source were available. PET was calculated using the Penman-Monteith equation (Guo et al. 2016). Based on the climatic water balance values a time-series of the Standardized Precipitation-Evapotranspiration Index (SPEI) was calculated (Vicente-Serrano et al. 2009). An important advantage of the SPEI is that it can be computed at different time scales. This way it is possible to incorporate the influence of the past values of the variable in the computation, and the magnitude of these values is by parameter scale. For example, a value of three would imply that data from the current month and of the past two months will be used for computing the SPEI or SPI value for a given month.

### 2.3 Population growth rate model

Following an analysis pipeline developed for the Glanville fritillary metapopulation (Kahilainen et al. 2018), we fitted a Bayesian linear mixed effects model to study the association between weather conditions and population growth rate at the regional scale. Instead of using monthly temperature and precipitation values, we used monthly water balance indices, which are more explicitly associated with drought conditions. We first divided individual habitat patches into groups of spatially clustered semi-independent habitat patch networks (SIN; see Hanski et al. 2017) and derived yearly population growth rates (*r*) as:

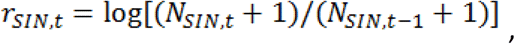

where *N*_*SIN,t*_ is the number of overwintering larval nest found in a SIN in the fall of year *t*. We then extracted water balance data for each SIN (see previous section) and fitted a linear mixed model for population growth rate using the water balance values across different life-history stages of the butterfly as covariates. In the model we included a random intercept and a first order autocorrelation term for each SIN. We implemented the model in Stan statistical modelling platform (Carpenter et al. 2017), using R packages brms (version 2.7.0; Bürkner 2017) and RStan (2.17.3; Stan Development Team 2018) as its interface. For further details regarding the analysis pipeline and implementation of the model, see Kahilainen et al. 2018. From the model we then obtained fitted annual growth rates for each of the 33 SINs included in the data and compared these with the observed growth rate values (Fig. 3).

### 2.4 Extinction probability model

We fitted a linear mixed effect model (logit link function) to binary patch extinction data calculated for each year of the survey between 1999 and 2018 to examine whether the high extinction rates in 2018 could be attributable to covariates previously recognized to impact the extinction probability of local populations (Hanski et al. 1995; Saccheri et al. 1998). Year was included as a random effect. Environmental variables included as fixed effects were the natural logarithm of patch area, the amount of the dominant host plant in each patch estimated on a scale of 1-3, the percent of host plants that were dried out, and the percent of the patch that was grazed (see Ojanen et al. 2013 for more details). Historical contingency was accounted for by considering the natural logarithm of the number of larval nests in each focal patch in the previous year, and the number of larval nests in the neighbourhood of each focal patch in the previous year, defined as:

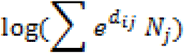

where, *d* is the distance between focal patch *i* and *j*, and *Nj* is the number of nests in patch *j* in the previous year. We trained the model using 80% of the data, and subsequently assessed model performance – quantified using the area under the receiver operating characteristic (AUC) – on the remaining 20% of the data. This procedure was repeated 10 times in order to assess the sensitivity of model performance to the train/test split.

## 3. RESULTS

### 3.1 Demographic declines across the butterfly metapopulation in 2018

Despite systematic surveying efforts, an all-time low of only 91 larval nests were recorded during the autumn survey in 2018 (the average number of nests recorded each year is 2750; figure 1). The number of occupied habitat patches was also an order of magnitude lower than in any average year (N=54; figure 1A), and only a single colonization event was recorded. The number of larval nests recorded within the occupied habitat patches was approximately 65% below the average (table S1).

**Figure 1.**
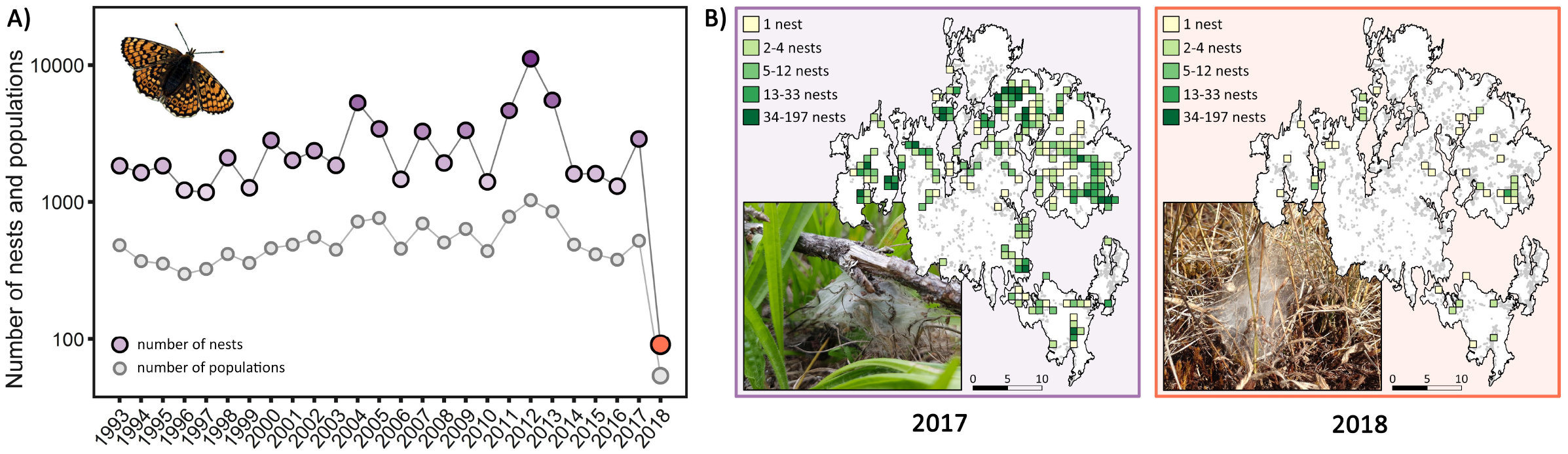
Population dynamics and demographic decline of the Glanville fritillary butterfly metapopulation on the Åland islands, Finland. A) Population dynamics of the butterfly metapopulation in years 1993-2018. The coloured dots represent the number of nests counted in yearly surveys, and the grey dots the number of occupied habitat patches in each year (y-axis represents a 10-log scale). Photograph depicts a female Glanville fritillary butterfly, Melitaea cinxia (photo credit: Tari Haahtela). B) The number oflarval nests found during autumn surveys on the Åland islands in 2017 (left-hand panel; average climatic conditions) and 2018 (right-hand panel; extreme climatic event). Coloured squares indicate the number of nests per square kilometre. Surveyed but unoccupied habitat patches are shown in grey. Photographs represent larval nests on a typical Plantago lanceolata host plant (left-hand panel; photo credit: Ana Salgado), and on P. lanceolata affected by drought (right-hand panel; photo credit: Felix Sperling).

### 3.2 Large deviations from past climatic conditions

Like many regions in Northern Europe, the habitat patches on the Åland islands received remarkably low levels of precipitation in the summer of 2018. The effect of this shortage was intensified by record-breaking temperatures (figure S1), resulting in the lowest climatic water balance values on record at the Jomala weather station in Åland (1960-2018; figure 2A). Consequently, the summer of 2018 was anomalous also in terms of primary productivity. Satellite-derived vegetation indices [Normalized Difference Vegetation Index (NDVI) and Enhanced Vegetation Index (EVI)], available since 2000, demonstrate low levels of vegetation productivity within the pastures and dry meadows inhabited by the butterfly in 2018 (figure 2B). The latter likely reflects also reduced host plant quality and availability during the periods when the post- and pre-diapause butterfly larvae are feeding (see figure 1B).

**Figure 2.**
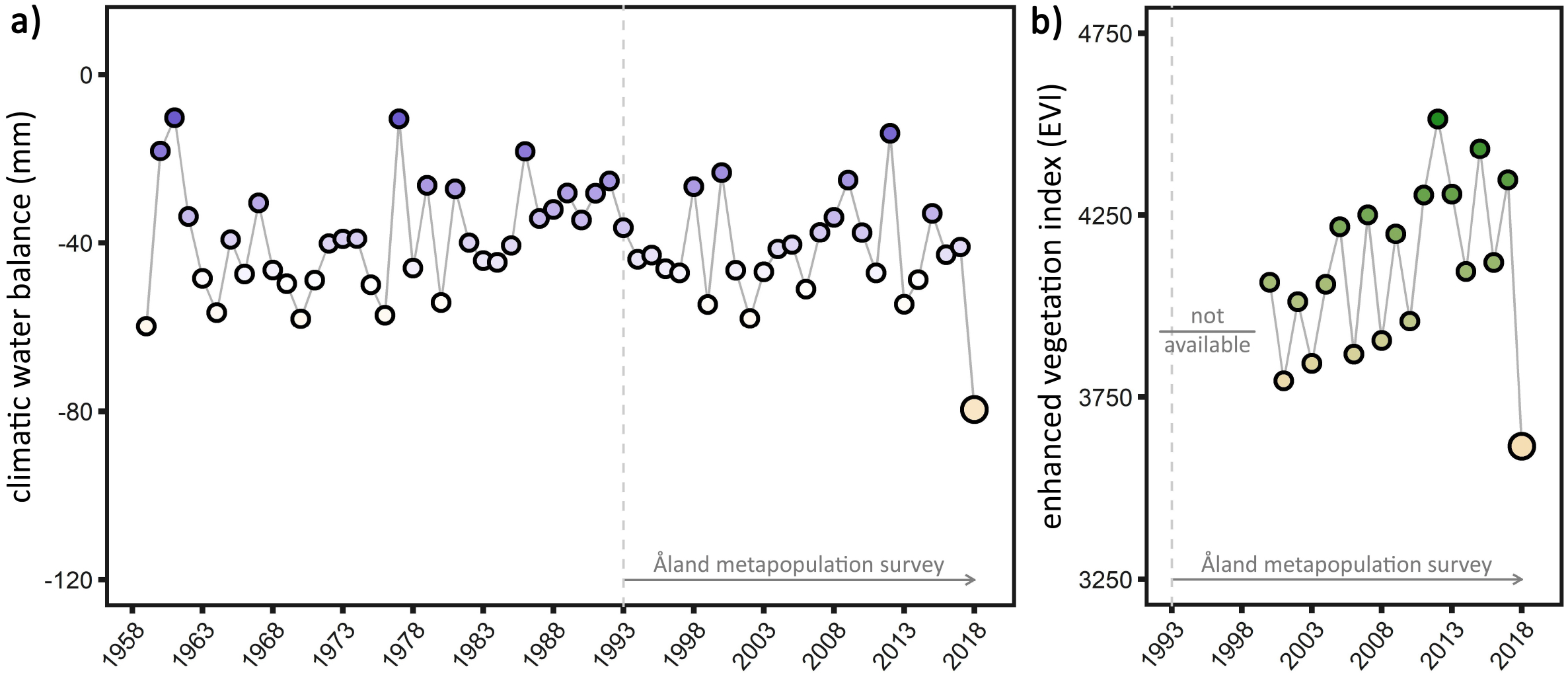
Meteorological and vegetation index data showing the extremity of the drought in 2018. A) Mean water balance (mm; precipitation - potential evapotranspiration) in April-August in years 1961-2018 derived from the Jomala climate station database in Åland. B) Mean Enhanced Vegetation Index (EVI; a measure of plant productivity) in the 4,415 habitat patches of the Glanville fritillary butterfly in the Åland islands in June-August from 2000 to 2018 derived from bands 1 and 2 of the MODIS on board NASA’s Terra satellite. All climatic variables and monthly values are represented in figure S1.

**Figure 3.**
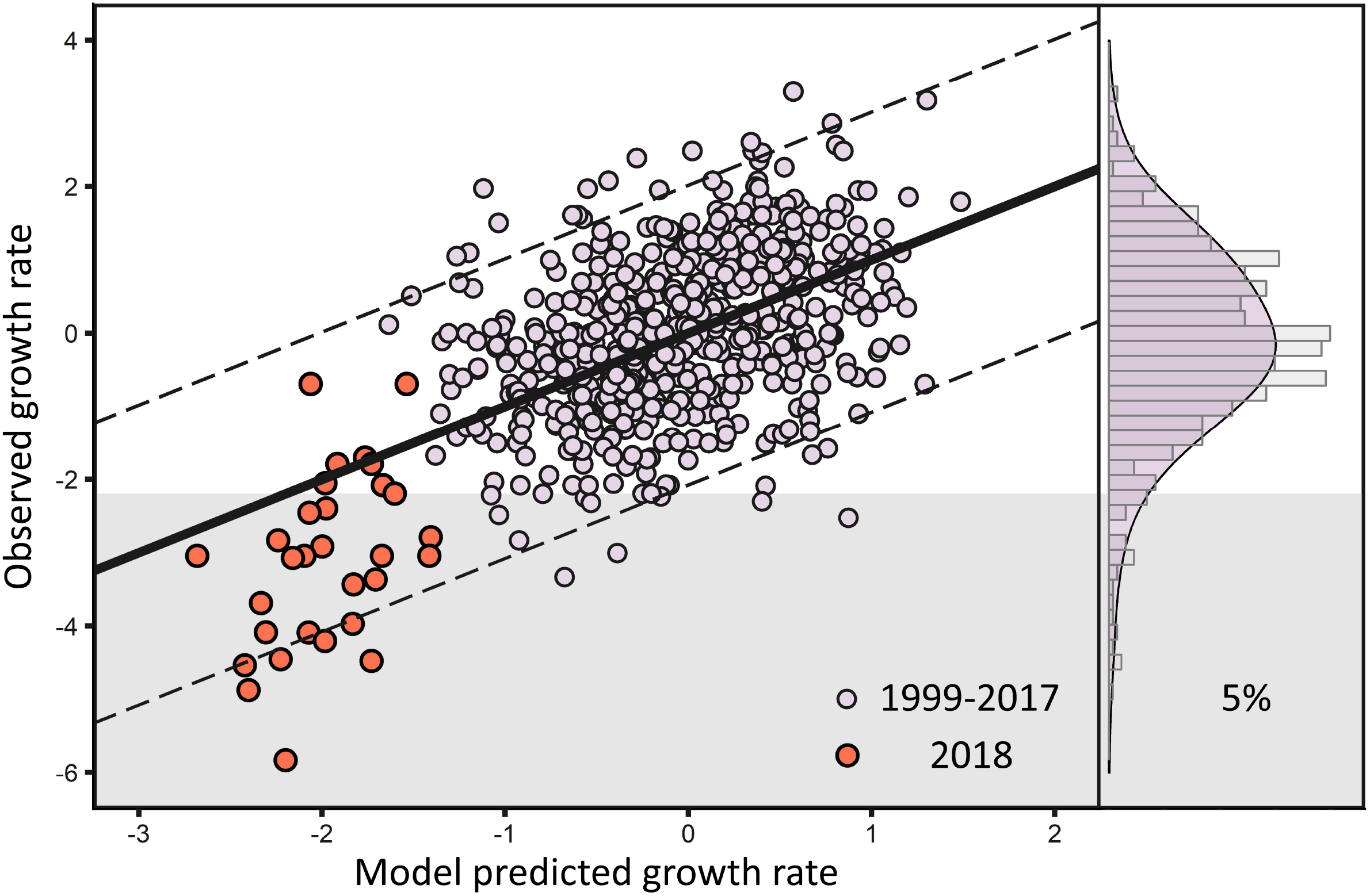
The relationship between observed and fitted population growth rates of the Glanville fritillary butterfly populations in the Åland islands from a linear mixed effects model with water balance values across the butterfly’s life cycle as predictors of population growth rate at the semi-independent network level. The purple dots illustrate observations for years 1999-2017, while observations for 2018 are given in red. Dashed lines represent the 95% interval of model residuals. The right-hand panel shows a histogram of the distribution of population growth rate values in 1999-2018, with the shaded area highlighting the values in the lowest 5% range of the population growth rate distribution.

### 3.3 Drought drives regional population declines

We examined whether the extreme climatic conditions experienced by the butterfly in 2018 were sufficient to drive population declines at the scale of the semi-independent patch networks (SINs; Hanski et al. 2017). Overall, monthly water balance indices – especially the water deficits during May and July – were highly associated with population declines at the SIN level (figure 3; table S2). With few exceptions, the model residuals for 2018 were negative, indicating that the observed declines in most SINs were more severe than suggested by the model fit (figure 3). Indeed, SIN level population declines in 2018 were among the most dramatic ones observed since the start of the annual survey, with nearly 80% of the growth rate values falling within the lowest 5 percentile (figure 3; table S1).

### 3.4 Summer drought decreases the predictability of local extinctions

We first confirmed the previously identified factors affecting extinction probability of local populations in this system. In short, the probability of extinction is lower in habitat patches that are large and well-connected. Patches with low population densities in the previous year, reduced host plant availability, or those used for livestock grazing, have higher extinction probabilities (table S3). Secondly, we estimated the predictive accuracy of our environmental extinction model. The mean AUC score across the 20 years was 0.76, implying that model covariates predict the extinction of a local population reasonably well (figure 4a; AUC scores above 0.80 reflect good discrimination (Swets 1988). The AUC scores varied significantly across years with the lowest values observed in years that are characterized by water deficits during summer (figure 4b). Model covariates were more likely to predict the extinction probability correctly in the years prior to the ECE of 2018.

**Figure 4.**
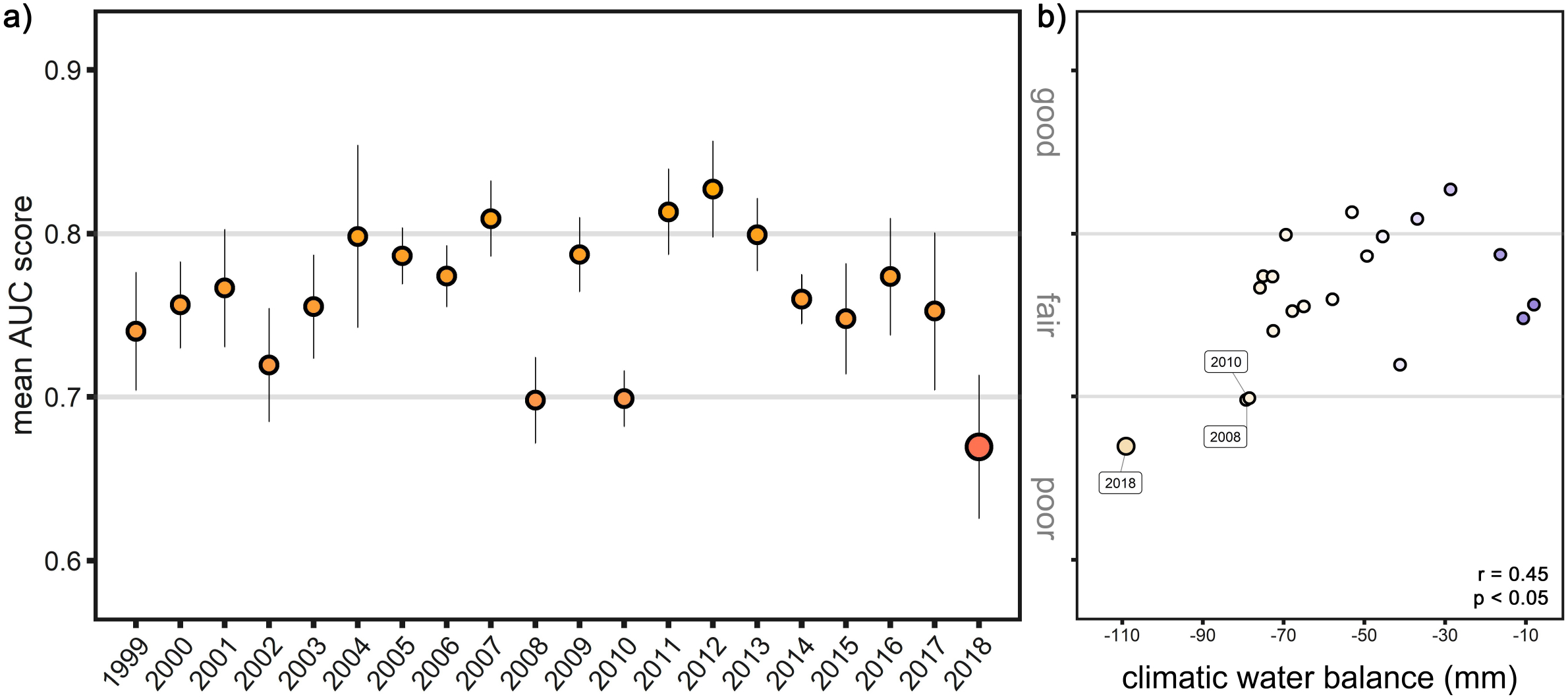
The predictability of local extinctions in a butterfly metapopulation. A) Model performance – quantified using the area under the ROC (receiver operating characteristic) curve – declined considerably in 2018, suggesting the extinction probability could not be accurately estimated using information on patch area, resource abundance, grazing, and population density in the previous year (see table S3). Meanwhile, these covariates captured extinction risk reasonably well in other years. Models were trained on ten random subsets of 80% of the available data. Plotted values correspond to mean values and line segments to standard error **bars**. As a general rule an AUC score of ≤ 0.7 is considered poor discrimination, 0.7 ≤ AUC score < 0.8 is considered fair, and 0.8 ≤ AUC score < 0.9 is considered good discrimination (Swets 1988). B) Some of the other years showing reduced model performance (e.g. 2008 and 2010) are also characterized by water deficits during summer. The x-axis represents the mean water balance (mm; precipitation - potential evapotranspiration) in May-July, and Spearman’s rho and p-value are displayed in the plot.

## 4. DISCUSSION

The ongoing climate crisis is expected to result in a rapid increase in the frequency, intensity and duration of extreme climatic events (e.g. Sun et al. 2014; Fischer & Knutti 2015). Owing to the profound impacts these extreme events can have on ecological functioning, the vulnerability of natural populations to the impacts of climate change may be severely underestimated (Anderson et al. 2017; Ummenhofer & Meehl 2017).

The Åland islands, like many other regions across the northern-hemisphere, sustained severe drought in the summer of 2018. Using extensive long-term data of the spatially-structured natural population of the Glanville fritillary butterfly, we demonstrate that both the climatic conditions during the summer and biological response of the metapopulation exceeded the 5% threshold values typically used to define ECEs (van de Pol et al. 2017). The majority of studies investigating the impacts of ECEs have focussed on the ability of organisms to cope with the direct effects of the extreme environmental conditions (Hoffmann & Sgrò 2018). While an organisms’ critical physiological limits are likely to be important risk factors, extinctions of local populations could also be triggered indirectly (Maron et al. 2015; Johansson et al. 2020). Our estimates of the occupancy and abundance of local populations across the archipelago are collected in the autumn, and hence only represent populations that survived the potentially harsh summer conditions. Therefore, we cannot infer whether the population declines and local extinctions were triggered directly by the arid conditions, or indirectly through for example climate-induced restrictions in resource availability (Maron et al. 2015). Nonetheless, using satellite-derived vegetation indices we demonstrate here that productivity was substantially reduced within the habitat patches of the butterflies in 2018. These data suggest that the severe drought negatively impacted the quality and availability of the host plants of these herbivorous insects, making resources scarce when demands were high. These results are in line with detailed observations made in 12 focal patches in 2018, where despite observing large numbers of clutches in early summer only two larval nests were found in autumn (Salgado et al. 2020).

We explored whether the strong deviations from usually experienced conditions influenced our ability to forecast the short-term biological response of the metapopulation, both at a regional and local scale. First, we demonstrate that the population growth rates at regional scales (i.e. the level of SINs; Hanski et al. 2017) are positively associated with climatic water balance during summer months (table S2). In other words, population sizes of this drought-sensitive butterfly species declined in years that were characterized by water deficits during summer (see also Oliver et al. 2015; Kahilainen et al. 2018). This result strongly points towards climatic conditions as the key driver of the dramatic decline of the metapopulation in 2018. In addition, we find that the relationship between the climatic variables and the biological response of the metapopulation is largely linear (i.e. model residuals of 2018 are negative, but generally do not exceed model confidence limits; figure 3; van de Pol et al. 2017). These results highlight that our detailed understanding of the population dynamics of this ecological system allowed us to – at least in part – forecast the observed population declines in response to the described extreme event.

While we were able to capture climate-induced changes in dynamics at regional scales, we were less successful in predicting the extinction probabilities of local populations in 2018 (figure 4a). In general, the model covariates, such as habitat size and connectivity, were likely to predict the extinction probability correctly in years when water availability was reduced during summer. In 2018, even populations inhabiting large and well-connected patches with high abundance in the previous year went extinct. This suggests that summer droughts, and in particular the extreme climatic conditions of 2018, may force the population over a threshold in which local extinctions are driven by factors other than those highlighted in previous studies with the system (reviewed in Ovaskainen & Saastamoinen 2018). Hence, uncharacterised ecological variables, such as the water holding capacity of the patch and/or the microhabitat heterogeneity within each meadow or pasture, may potentially be important determinants of population persistence under arid environmental conditions. In addition, stochastic processes may become more important in dry years.

As a potential caveat we note that, despite using a systematic and methodical protocol, our probability to detect the presence of the butterfly is known to be reduced in very small populations (Hanski et al. 2017). Hence, strong climate-induced declines in population sizes may therefore contribute to decreased detection rates (and, consequently, an overestimation of the biological response). Other variables, such as reduced nest sizes or shifts in habitat choice, may potentially also negatively affect detectability under climatic extremes. However, our surveying protocol dictates that when a patch is judged to be unoccupied, it should immediately be re-surveyed with the same effort (Ojanen et al. 2013). Owing to this safeguard mechanism the majority of the patches were surveyed twice in 2018, reducing the potential influence of the ECE on nest detectability.

The documentation of the demographic crash of the Glanville fritillary metapopulation highlights the vulnerability of natural populations and underscores the importance of long-term monitoring programs to effectively document the consequences of ECEs on species, population and community dynamics. Our results further demonstrate that ECEs can impede conservation efforts by reducing the value of predictive models. For example, even within this extensively understood study system any effort to potentially mitigate the impacts of the ECE on local populations would likely have failed, since we would not have been able to predict which local populations were most vulnerable. The long-term implications of the ECE of 2018 are yet to be realized and continuation of the metapopulation survey during the following years will demonstrate whether the disruption from the normal extinction-colonization dynamics has driven this iconic metapopulation close to its tipping point (Dai et al. 2013) or whether the remaining populations will recover from this rare perturbation event. In the case of the latter, this will provide a unique and exciting opportunity to study the long-term ecological and evolutionary impacts of an ECE in a spatially-structured population.

## Acknowledgements

We are grateful to all researchers and field assistants that have participated in the field work on Åland between 1993 and 2018. Financial support was provided by the European Research Council (StG grant META-STRESS no. 637412 to M.S.). The authors declare no conflicts of interest.

## Authors contributions

All authors contributed to the conception and design of the work. MS and ML contributed to the acquisition of the data. TD, MFD and AK performed the analyses and EvB and AM prepared the figures. EvB, AM and MS wrote the first draft of the manuscript. All authors contributed and approved the final manuscript.

## Data availability

Datasets used in this study are available online at (will be deposited in Dryad once accepted for publication).

## Supplementary Materials

Tables S1 - S3; Figure S1

---

**Supplementary Table 1.** Overview of the Glanville fritillary metapopulation long-term data and population growth rates in the viable semi-independent habitat patch networks (SINs).

| Long term data |  |  |  |  |  | SIN growth rates |  |
| --- | --- | --- | --- | --- | --- | --- | --- |
| Year | Patches | Populations | Nests | %Occupied | Abundance | GR<sub>SIN</sub> (N) | % in lower 5% |
| 1993 | 1324 | 485 | 1834 | 36.63 | 3.781 | na | na |
| 1994 | 1379 | 372 | 1636 | 26.98 | 4.398 | 33 | 12.1 |
| 1995 | 1374 | 356 | 1833 | 25.91 | 5.149 | 31 | 0.0 |
| 1996 | 1333 | 300 | 1218 | 22.51 | 4.060 | 31 | 3.2 |
| 1997 | 1507 | 326 | 1176 | 21.63 | 3.607 | 31 | 3.2 |
| 1998 | 2792 | 418 | 2101 | 14.97 | 5.026 | 32 | 0.0 |
| 1999 | 3927 | 360 | 1267 | 9.17 | 3.519 | 32 | 3.1 |
| 2000 | 2935 | 461 | 2808 | 15.71 | 6.091 | 33 | 0.0 |
| 2001 | 3951 | 490 | 2011 | 12.40 | 4.104 | 31 | 3.2 |
| 2002 | 3160 | 556 | 2359 | 17.59 | 4.243 | 32 | 0.0 |
| 2003 | 3160 | 450 | 1849 | 14.24 | 4.109 | 31 | 0.0 |
| 2004 | 3195 | 720 | 5308 | 22.54 | 7.372 | 32 | 0.0 |
| 2005 | 3881 | 765 | 3399 | 19.71 | 4.443 | 33 | 3.0 |
| 2006 | 3453 | 458 | 1461 | 13.26 | 3.190 | 33 | 6.1 |
| 2007 | 3693 | 698 | 3259 | 18.90 | 4.669 | 32 | 0.0 |
| 2008 | 3644 | 508 | 1922 | 13.94 | 3.783 | 31 | 3.2 |
| 2009 | 3905 | 638 | 3318 | 16.34 | 5.201 | 32 | 0.0 |
| 2010 | 3588 | 440 | 1401 | 12.26 | 3.184 | 32 | 15.6 |
| 2011 | 3796 | 783 | 4644 | 20.63 | 5.931 | 32 | 0.0 |
| 2012 | 3845 | 1031 | 11129 | 26.81 | 10.794 | 33 | 0.0 |
| 2013 | 3704 | 854 | 5540 | 23.06 | 6.487 | 32 | 0.0 |
| 2014 | 4371 | 490 | 1603 | 11.21 | 3.271 | 32 | 0.0 |
| 2015 | 4186 | 416 | 1608 | 9.94 | 3.865 | 33 | 3.0 |
| 2016 | 4203 | 381 | 1303 | 9.06 | 3.420 | 33 | 0.0 |
| 2017 | 4352 | 523 | 2871 | 12.02 | 5.489 | 30 | 0.0 |
| 2018 | 3343 | 54 | 91 | 1.62 | 1.685 | 30 | 76.7 |

Patches: the number of habitat patches included in the survey (survey extended between 1997-1999; peripheral islands not included in 2018); Populations: the number of patches that were occupied by the butterfly; Abundance: the mean number of nests per population; GR<sub>SIN</sub> (N) the number of viable SINs (33) for which a population growth rate could be estimated; % in lower 5%: percentage of population growth rates that fall within the 5% most negative values recorded in the Åland islands (see figure 2 in the main text).

**Supplementary Table 2.** Estimated coefficients, their estimated standard errors, and 95% credible intervals for the model estimating the association between the standardized climatic water balance and SIN growth rates.

| Covariate | Est. coef. | Est. SE | 95% Cr.I. |  |
| --- | --- | --- | --- | --- |
|  |  |  | Lower | Upper |
| Intercept | -0.1105 | 0.0324 | -0.1745 | -0.0467 |
| WaB<sub>Diapause</sub> | -0.1247 | 0.0463 | -0.2159 | -0.0327 |
| WaB<sub>Mar</sub> | -0.0874 | 0.0350 | -0.1562 | -0.0184 |
| WaB<sub>Apr</sub> | 0.1384 | 0.0468 | 0.0470 | 0.2295 |
| WaB<sub>May</sub> | 0.3030 | 0.0518 | 0.2008 | 0.4033 |
| WaB<sub>Jun</sub> | 0.2339 | 0.0428 | 0.1493 | 0.3178 |
| WaB<sub>Jul</sub> | 0.3480 | 0.0541 | 0.2404 | 0.4535 |
| WaB<sub>Aug</sub> | 0.1361 | 0.0393 | 0.0586 | 0.2130 |
| AR[1] | -0.2913 | 0.0391 | -0.3678 | -0.2152 |
| $\sigma_{(\text{SIN intercept})}$ | 0.0714 | 0.0462 | 0.0032 | 0.1713 |
| $\sigma_{\text{res}}$ | 1.0304 | 0.0260 | 0.9804 | 1.0821 |

AR[1]: first-order autocorrelation term; WaB<sub>Diapause</sub>: Average water balance during the diapause period; WaB<sub>Mon</sub>: Monthly water balance; $\sigma_{(\text{SIN intercept})}$ : standard deviation of random intercepts; $\sigma_{\text{res}}$ : residual standard deviation.

**Supplementary Table 3.**
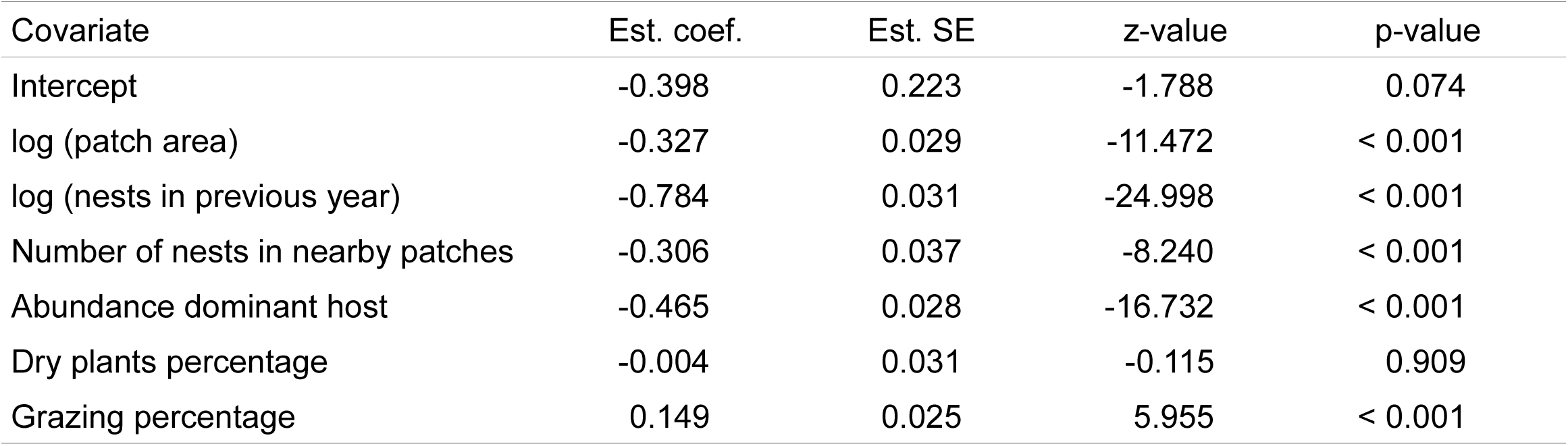
Coefficients from a fit linear mixed effects model with year as a random effect, highlighting the importance of patch area, resource abundance, grazing, and population demography from the previous year on extinction probability. Data include 9784 potential extinction events in the last two decades.

| Covariate | Est. coef. | Est. SE | z-value | p-value |
| --- | --- | --- | --- | --- |
| Intercept | -0.398 | 0.223 | -1.788 | 0.074 |
| log (patch area) | -0.327 | 0.029 | -11.472 | < 0.001 |
| log (nests in previous year) | -0.784 | 0.031 | -24.998 | < 0.001 |
| Number of nests in nearby patches | -0.306 | 0.037 | -8.240 | < 0.001 |
| Abundance dominant host | -0.465 | 0.028 | -16.732 | < 0.001 |
| Dry plants percentage | -0.004 | 0.031 | -0.115 | 0.909 |
| Grazing percentage | 0.149 | 0.025 | 5.955 | < 0.001 |

**Supplementary Figure 1.**
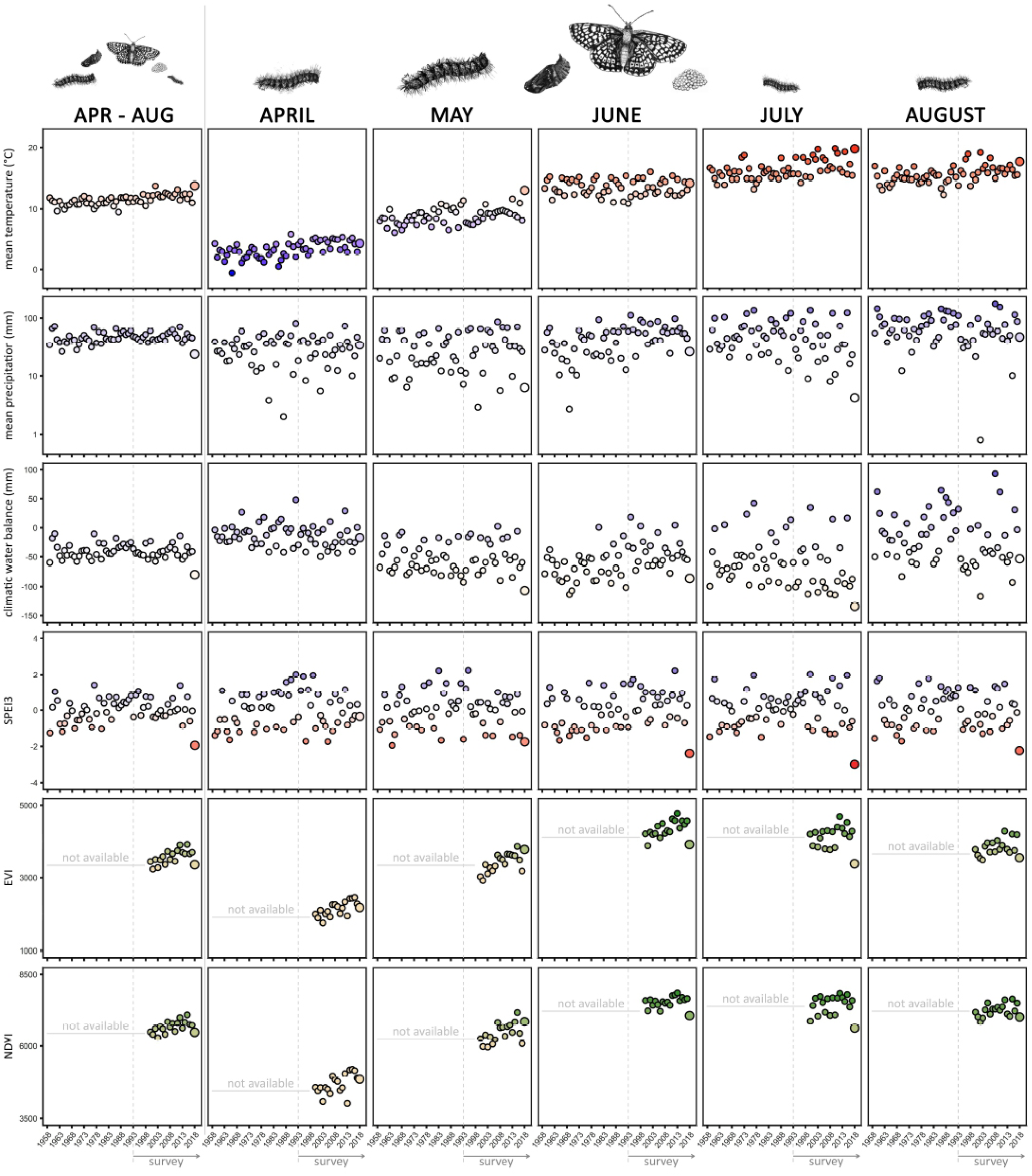
Meteorological (Mean temperature; Mean precipitation; Mean climatic water balance (mm; precipitation - potential evaporation); SPEI3: Mean Standardised Precipitation-Evapotranspiration Index at a 3 month scalar) and vegetation indices (EVI: Mean Enhanced Vegetation Index; NDVI: Mean Normalized Difference Vegetation Index) data for the period of April-August (left), and then separately for each month corresponding to the different life stages of the butterfly. The meteorological data in 1958-2018 are derived from the Jomala climate station database in Åland. The vegetation index data were derived from bands 1 and 2 of the MODIS on board NASA’s Terra satellite and calculated for the 4,415 habitat patches of the Glanville fritillary butterfly in the Åland islands. Illustrations courtesy of Luisa Woestmann.

